# Towards an archaeological workflow for sedaDNA sample collection: methods and best practices for minimizing surface contamination

**DOI:** 10.1101/2025.09.01.673420

**Authors:** Kadir Toykan Özdoğan, Fabricio Furni, Ivo Laros, Gertjan Plets, G. Arjen de Groot

**Author notes:** Corresponding author: Kadir Toykan Özdoğan.

## Abstract

SedaDNA has been successfully recovered from diverse contexts, enabling researchers to address paleoenvironmental questions such as ecosystem changes and climatic shifts, along with archaeological questions related to past human activities, diet, health, and interactions with their environment. Despite its potential, the reliability of sedaDNA data heavily depends on proper sampling and handling practices. Contamination remains a significant challenge in ancient DNA research, profoundly impacting data integrity and interpretation. Consequently, numerous studies have underscored the importance of minimizing contamination levels. Nonetheless, a challenge arises in implementing this in daily archaeological practice, as a specialized DNA expert is not always available. This study suggests two customised workflows of commonly used sedaDNA sampling techniques, redesigned for archaeologists without sedaDNA expertise. We test their effectiveness in the field. To benchmark these protocols, we applied artificial DNA contaminants onto sediment surfaces before sampling. Subsequently, we evaluate archaeologists’ effectiveness in reducing surface contamination by quantifying the level of artificial DNA detected. Finally, we compare results between samples taken by archaeologists with and without sedaDNA expertise. Our results demonstrate that existing sampling protocols, when slightly adapted to the needs of archaeologists, significantly minimize surface contamination to the point that they provide high-quality extraction. Additionally, our findings suggest that subsampling sediments with minimal surface contamination from previously collected materials is viable, emphasizing the potential of archived sediment collections for sedaDNA analysis.

## Introduction

Ancient DNA recovered in archaeological sediments (sedaDNA) is a powerful tool for reconstructing past ecosystems and studying biodiversity changes (Alsos et al., 2024; Nguyen et al., 2023). Different projects have shown that extracting ancient DNA from sediments allows for detecting organisms consumed or living near a site (Aldeias & Stahlschmidt, 2024). A sedaDNA approach offers unique insights into environmental and archaeological questions, helping researchers reconstruct the past with distinctive detail.

Over the last decade, sedaDNA has been successfully recovered from diverse contexts, ranging from lake and marine sediments to permafrost, swamp forests, and rock shelters. These studies have primarily addressed paleoenvironmental questions like ecosystem changes and climatic shifts (L. Armbrecht et al., 2022; Braadbaart et al., 2020; Crump et al., 2021; Dommain et al., 2020; Hebda et al., 2022; Kjær et al., 2022; Murchie et al., 2021, 2022; Pedersen et al., 2021). Meanwhile, sedaDNA from human-associated contexts like latrines, wells, ponds, middens, and cave sediments increasingly allows researchers to explore archaeological questions, focusing on past human activities, diet, health, and interactions with their environment (Boivin et al., 2024; Borry et al., 2020; Gelabert et al., 2021; Sabin et al., 2020; Seersholm et al., 2016; Søe et al., 2018; Vernot et al., 2021; Zavala et al., 2021). However, studies so far have only scratched the surface of the potential that sedaDNA has to offer for archaeology (Özdoğan et al., 2024).

One critical advantage of sedaDNA lies in its ability to recover genetic material from sediments even when no visible organic remains are present. This capability enables researchers to access a broader spectrum of biological data, including traces of plants, animals, and microorganisms that once inhabited or passed through an archaeological site (Aldeias & Stahlschmidt, 2024). Furthermore, Kjær et al. showed sedaDNA’s potential to recover ancient DNA from exceptionally old contexts (2022). Additionally, sedaDNA enhances the temporal and spatial resolution of focused regions, allowing scientists to track changes in biodiversity and ecosystems over time and space (Wang et al., 2021).

Despite its potential, the reliability of sedaDNA data depends heavily on proper sampling and handling practices. Contamination is a significant challenge in ancient DNA research, with profound implications for data integrity and interpretation (Orlando et al., 2021). Numerous studies have emphasized the importance of minimizing contamination levels, but it remains a persistent issue that researchers must carefully address (Fulton & Shapiro, 2019; Furtwängler et al., 2018; Llamas et al., 2017; Renaud et al., 2015; Skoglund et al., 2014).

Contamination in sedaDNA studies can broadly be categorized into two types: modern contamination and ancient cross-contamination. Modern contamination is typically introduced during excavation or sample handling. For instance, DNA from researchers, animals, or plants that are present at the site can mix with the ancient material (Ariza et al., 2023; Edwards et al., 2018; Peyrégne & Prüfer, 2020). While bioinformatic tools can filter out some modern contamination, such DNA is often abundant and can overwhelm authentic ancient signals (Michelsen et al., 2022; Peyrégne & Peter, 2020). This is especially problematic when modern and ancient sequences come from closely related taxa, complicating data interpretation (e.g., taxon identification) and potentially leading to incorrect conclusions (Orlando et al., 2021).

Ancient cross-contamination presents an even more significant challenge. Unlike modern contaminants, ancient contaminant DNA fragments share similar damage profiles with authentic ancient DNA, making them difficult to distinguish (Dabney et al., 2013). This type of contamination often occurs during excavation, storage, or laboratory processing. For example, disturbed sediments from different archaeological contexts may mix during excavation. Since ancient cross-contamination is challenging to identify and isolate, it significantly complicates the reliable reconstruction of the authentic ancient DNA signals in a contaminated sample (Llamas et al., 2017). Ancient DNA fragments from one sample can contaminate another in a shared storage facility or during laboratory processing. Using sterile tools, wearing protective gear, and isolating samples during collection and storage is critical (Knapp et al., 2012).

Archaeological excavations are logistically complex and fast-paced, often hindering the implementation of precise and controlled sediment sampling. Consequently, the risk of cross-contamination from sections’ surfaces increases when clean sampling protocols are not adhered to. Previous studies have utilized tracers to monitor surface contamination, thereby validating the reliability of samples (Epp et al., 2019; Wang et al., 2021). However, adopting careful protocols can significantly reduce contamination risks (Armbrecht et al., 2019), although contamination-free sampling may be impossible in most field conditions. These measures can help minimize contamination to acceptable levels, where bioinformatic filters are applied to remove taxa with lower assigned DNA reads and ancient-DNA-specific damage patterns (Michelsen et al., 2022).

While previous studies have explored the contamination risks in ancient DNA research (Dabney et al., 2013; Michelsen et al., 2022; Peyrégne & Peter, 2020), there is a limited understanding of how effectively non-experts can implement clean sampling protocols in their workflows. This is particularly relevant given the collaborative nature of archaeological projects, where teams with varying expertise often conduct sampling.

This study demonstrates that clean sampling can be effectively integrated into archaeological workflows. In doing so, we aim to facilitate the broader application of sedaDNA analysis. Enabling non-expert sampling will help researchers increase spatial and temporal coverage of their sample collections, eventually enhancing our ability to contribute to our understanding of past anthropogenic environments. To investigate this, we examined whether archaeologists with no prior aDNA expertise can successfully perform clean sampling protocols after receiving customized protocols and a tailored sampling kit supplemented with visual demonstrations. Prior to each trial, we deliberately sprayed a known amount of artificial DNA contaminants onto the sediment surfaces, allowing us to benchmark our workflows under known ‘worst-case’ conditions. Subsequently, we assessed the effectiveness of these methods in reducing surface contamination by quantifying the amount of artificial DNA in the collected samples. In addition to testing clean sampling methods in an excavation setting, we explored another approach to expanding sedaDNA applications: using previously collected pollen cores as sedaDNA sample sources. We analyzed surface contamination levels on previously collected cores while applying clean sampling protocols to determine the optimal subsampling locations. Additionally, we aimed to identify potential contamination sources and develop strategies to minimize them.

Through these different experimental scenarios, we highlight the potential of non-expert sedaDNA sampling when good practices are applied, both *in situ* and from previously collected sediment cores. In our discussion, we explore the implications of our findings for sedaDNA research in archaeology and present recommendations for integrating good sedaDNA sampling practices into archaeological workflows.

## Materials & Methods

### Experimental Design

Various methods have been used for sediment sampling for sedaDNA studies, reflecting the diverse contexts in which ancient DNA is preserved. Sediment samples have been collected directly from archaeological excavation sites, pollen cores, and even undisturbed archaeological remains such as pottery and bodily ornaments (Essel et al., 2023; Foley et al., 2012; Green & Speller, 2017; Murchie et al., 2022).

For this study, we focused on two primary sampling methods: (1) point-based sampling directly from open sections using sterile tubes (Wang et al., 2021) and (2) subsampling from the pollen core, which focuses on isolating uncontaminated core material by removing outer layers exposed to or in contact with potential contaminants using sterile tools. (Epp et al., 2019)

These two sampling approaches were selected because they are widely used in the field of sedaDNA and have the potential to highlight the specific challenges of surface contamination (Epp et al., 2019; Wang et al., 2021). Previous studies (Armbrecht et al., 2019; Seeber et al., 2024; Selway et al., 2022; Zimmermann et al., 2021) have demonstrated their effectiveness but relied on the involvement of experienced samplers. Two different protocols based on the above-mentioned sampling approaches were scripted and given to non-aDNA expert archaeologists (A.1, including a video instruction) for the written protocols and Data Availability for both written protocols and the video demonstration).

To assess the extent of surface contamination in sediment samples, we used synthetic double-stranded DNA fragments (gBlocks) as a contamination marker. In genetics research, gBlocks are widely utilized as PCR controls (Conte et al., 2018). Since any DNA sequence can be synthesized as gBlock DNA, this allowed us to use an artificial DNA sequence that is non-existent in nature (Appendix A.2) (Kavlick, 2018; Swango et al., 2006). Less than 24 hours prior to the experiment, we prepared solutions of gBlock DNA in 96% ethanol according to a protocol by Epp et al. (2019). Per artificial contamination event, we sprayed 100 µl of a solution with a concentration of 1e+7 copies per µl of gBlock molecules (A.2). This concentration was based on a pilot experiment to ensure that positive controls showed a positive result well within the detection range of the ddPCR platform.

During the experiment, gBlock DNA solutions were sprayed onto the sediment surfaces of the designated sampling points at an outdoor excavation site, the surfaces of the empty pollen core, and the sediment surface of the sampled pollen core. These points were carefully selected to represent realistic contamination scenarios. The selected clean sampling protocols (A.1) were then employed to collect sediment samples, with particular attention paid to removing contaminated surface layers and isolating uncontaminated material. Additional sediment samples were taken directly from the contaminated surface for use as positive controls.

To assess the effectiveness of the protocols mentioned above, residual levels of gBlock DNA in the collected sediment samples were quantified in the lab employing digital droplet PCR (ddPCR; see 3.3). By comparing the gBlock concentrations detected in experimental and control samples, we evaluated the effectiveness of the sampling protocols in minimizing surface contamination.

### Sampling

#### Point-based Sampling

Since previous studies have shown that soil types affect the preservation, absorption, and extraction of aDNA, we chose two different soil types for excavation sampling (Andersen et al., 2012a; Giguet-Covex et al., 2023). Sampling from clay-rich contexts was performed at an archaeological site in Tiel, the Netherlands, and sampling from sand-rich contexts was conducted at an archaeological site in Loenen, the Netherlands.

The protocol used for the point-based sampling, as described in detail in A.1, includes the following basic steps:

- Protective Gear: Researchers wore face masks and gloves to prevent contamination caused by them. Gloves were sterilized before sampling with bleach and replaced after each sample collection.
- Sterile Tools: Sterile spatulas were used to remove the surface layer of the section.
- Sampling Process: After clearing the surface, sterile centrifuge tubes were used to collect sediment. The tubes were pushed into the sediment profile and withdrawn with the filled sediment.
- Storage and Labeling: Samples were placed in zip-lock bags. Sticker labels were used to mark each sample.

Prior to the sampling, the non-expert participants received a copy of the protocol and brief visual demonstrations to familiarize themselves with the workflow and basic contamination-minimizing strategies. Before sampling, gBlock DNA solutions were sprayed onto the vertical archaeological section by the first author at three separate locations per participant, adding a control location. These locations were carefully marked to ensure consistency in sampling conditions.

First sediment samples were taken from the surface of the sprayed control location as positive controls. Then, the participants collected sediment samples from the specified points following the clean sampling protocol (Fig. 1 for detailed methodology). The participants included one sedaDNA expert and seven non-expert archaeologists (three at the clay-rich site, four at the sand-rich site) with field experience but no sedaDNA sampling experience. After collection, the sediment samples were securely stored in zipped bags at 4°C for three days before the DNA extractions at the Laboratory for Ecological Genetics of Wageningen Environmental Research (Wageningen, The Netherlands) (WENR).

**Figure 1.**
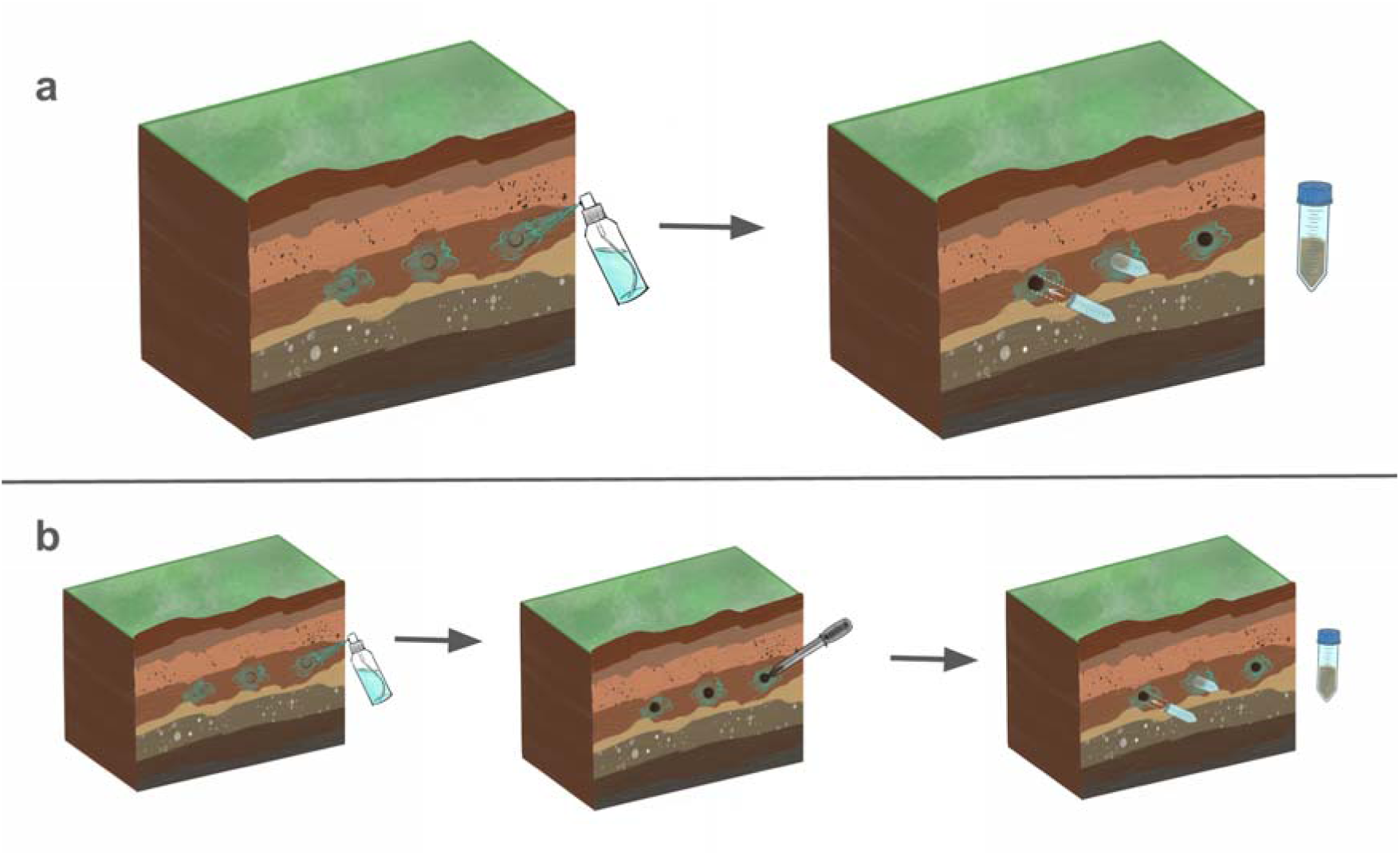
Point-Based Sampling Procedure: The first row (a) demonstrates the collection of surface samples: (1) Spraying the section with gBlock DNA solution, (2) Collecting contaminated surface soil using sterile tubes by inserting them into cleaned areas. The second row (b) outlines the collection of clean samples: (1) Spraying the section with gBlock DNA solution, (2) Cleaning artificially contaminated areas using sterile spatulas, (3) Collecting soil from the uncontaminated section using sterile tubes by inserting them into the cleaned areas.

#### Pollen Core Sampling

The protocol used for pollen core subsampling for sedaDNA is described in detail in A.1. It required the following basic steps:

- Protective Gear: Face masks and gloves were used to prevent contamination. Gloves were sterilized with bleach before subsampling and replaced after each subsample collection.
- Sterile Tools: Sterile spatulas were used to remove the surface layer of the core.
- Sampling Process: After cleaning the surface, sterile spatulas were used to fill the sterile centrifuge tubes with sediment.
- Storage and Labeling: Samples were placed in zip-lock bags. Sticker labels were used to mark each sample.

Pollen core sampling was conducted at the Tiel archaeological site, focusing on replicating conditions that might occur during sediment sampling from pollen cores in the field. The procedure began with applying gBlock DNA solutions to the inner surfaces of the pollen core. Furthermore, as for the point samples in 3.2.1, the vertical section where the coring was conducted was also sprayed with the gBlock DNA solution to simulate artificial surface contamination from every angle. Once the sampling was completed, the pollen core was transported to the WENR for subsampling.

Before subsampling, the surface of the sediment sample in the pollen core was also sprayed with the gBlock DNA solution. Subsampling was conducted in a controlled but non-sterile room, replicating the conditions typically found in archaeological storage facilities or rooms that do not meet the strict standards of ancient DNA laboratories. This setup aimed to evaluate the impact of such conditions on contamination control. Six subsamples were collected from the core: two from the surface, two from the midpoint, and two from the bottom of the core (Fig. 2).

**Figure 2.**
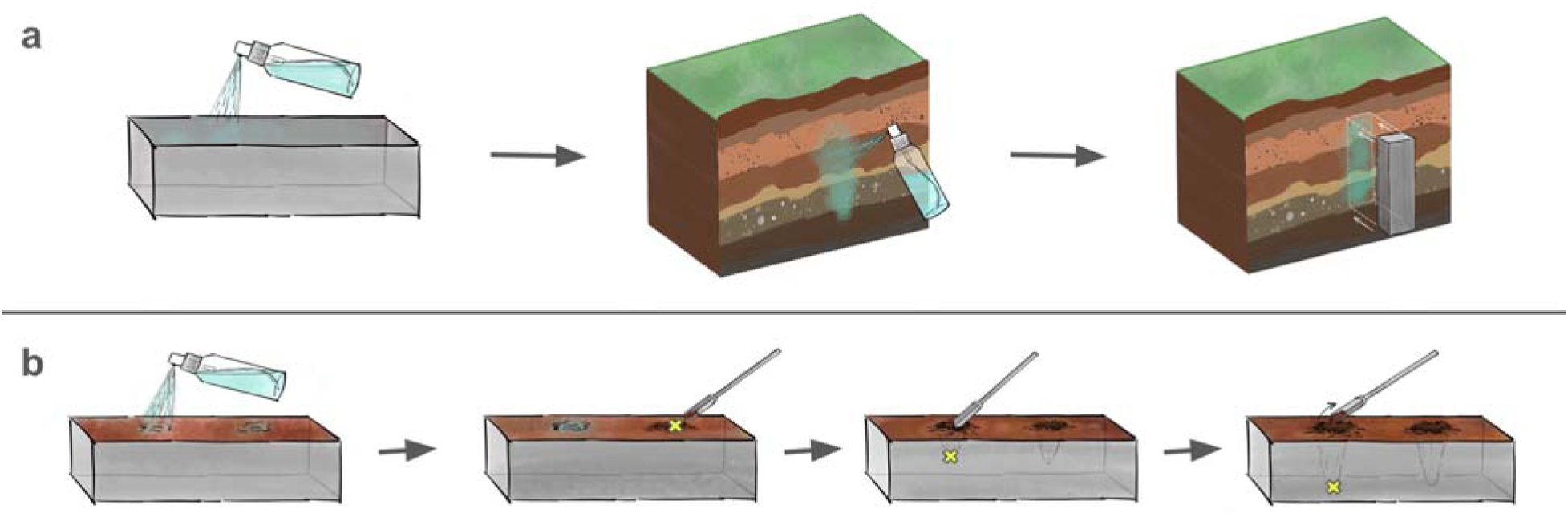
Pollen Core Sampling Procedure: The first row (a) illustrates the steps at the excavation site: (1) Spraying the empty pollen core with gBlock DNA solution, (2) Spraying the marked section of the core with gBlock DNA solution, (c) Inserting the pollen core to collect a layered soil sample. The second row (b) details the subsampling steps performed in a controlled environment: (1) Spraying the surface of the soil sample within the pollen core with gBlock DNA solution, (2) Collecting surface samples, (3) Sampling from the midpoint of the pollen core, (4) Sampling from the bottom of the pollen core.

#### Laboratory and Data Analysis

After collection, the sediment samples were taken to a dedicated DNA extraction laboratory for processing and further analysis. This phase aimed to assess the residual levels of gBlock DNA contamination in samples and evaluate the effectiveness of the sampling protocols in minimizing surface contamination. DNA extractions were performed in a dedicated, clean laboratory environment at WENR. The *DNeasy PowerSoil Pro Kit (QIAGEN)* was used for DNA extraction, following the manufacturer’s protocol. This kit was chosen due to its high efficiency in isolating DNA from complex and challenging sediment matrices, making it particularly suited for sedaDNA studies (*DNeasy PowerSoil Pro Kit Handbook – QIAGEN*, n.d.).

After extraction, the DNA samples underwent quantitative analysis using ddPCR. This technique uses a water-oil emulsion to fractionate a DNA extract in ∼20000 droplets. An isolated PCR amplification takes place in each droplet containing a target DNA molecule, after which fluorescence-based detection is used to count the number of droplets containing a target DNA molecule. Via this approach, ddPCR is capable of measuring contamination levels with high sensitivity and precision (Hunter et al., 2017; Wood et al., 2019). By providing an absolute count of DNA molecules, ddPCR is well suited for detecting even low-level contamination and quantifying the presence of gBlock DNA markers in the samples (Taylor et al., 2017).

Duplicate ddPCR reactions were performed for each DNA extract. A detailed description of the laboratory protocol applied for ddPCR analysis is provided in A.2. Raw output data of the ddPCR analysis were analyzed using BioRad QuantaSoft version 1.7.4.0817. For each replicate emulsion reaction, raw data represent the total number of droplets and the number of positive droplets (i.e., in which target DNA was amplified, resulting in a fluorescent signal). Because positive droplets may incidentally contain >1 target DNA molecule, for each reaction, the software applies Poisson distribution analysis to estimate the concentration of target DNA molecules in the template. Subsequently, these estimated concentrations per replicate reaction were used for further statistical data analysis.

After obtaining the quantitative results from ddPCR, we assessed the statistical significance of contamination values using pairwise two-tailed Student T-tests between sampling types and points. Specifically, we compared surface values from the excavation with those from the participants and separately analyzed the different sampling points within the pollen core.

## Results

### Sampling at an excavation

#### Clay-rich soil

In total, fifteen sediment samples were collected: (a) three artificially contaminated surface sediment samples, (b) three samples collected by an aDNA expert, and (c) nine samples collected by three non-expert participants (three samples each).

The surface sediment samples showed high levels of artificial gBlock DNA contamination, confirming that the experimental setup effectively simulated surface contamination (Fig. 3). This provided a robust baseline for evaluating the effectiveness of the clean sampling protocols performed by every participant.

**Figure 3:**
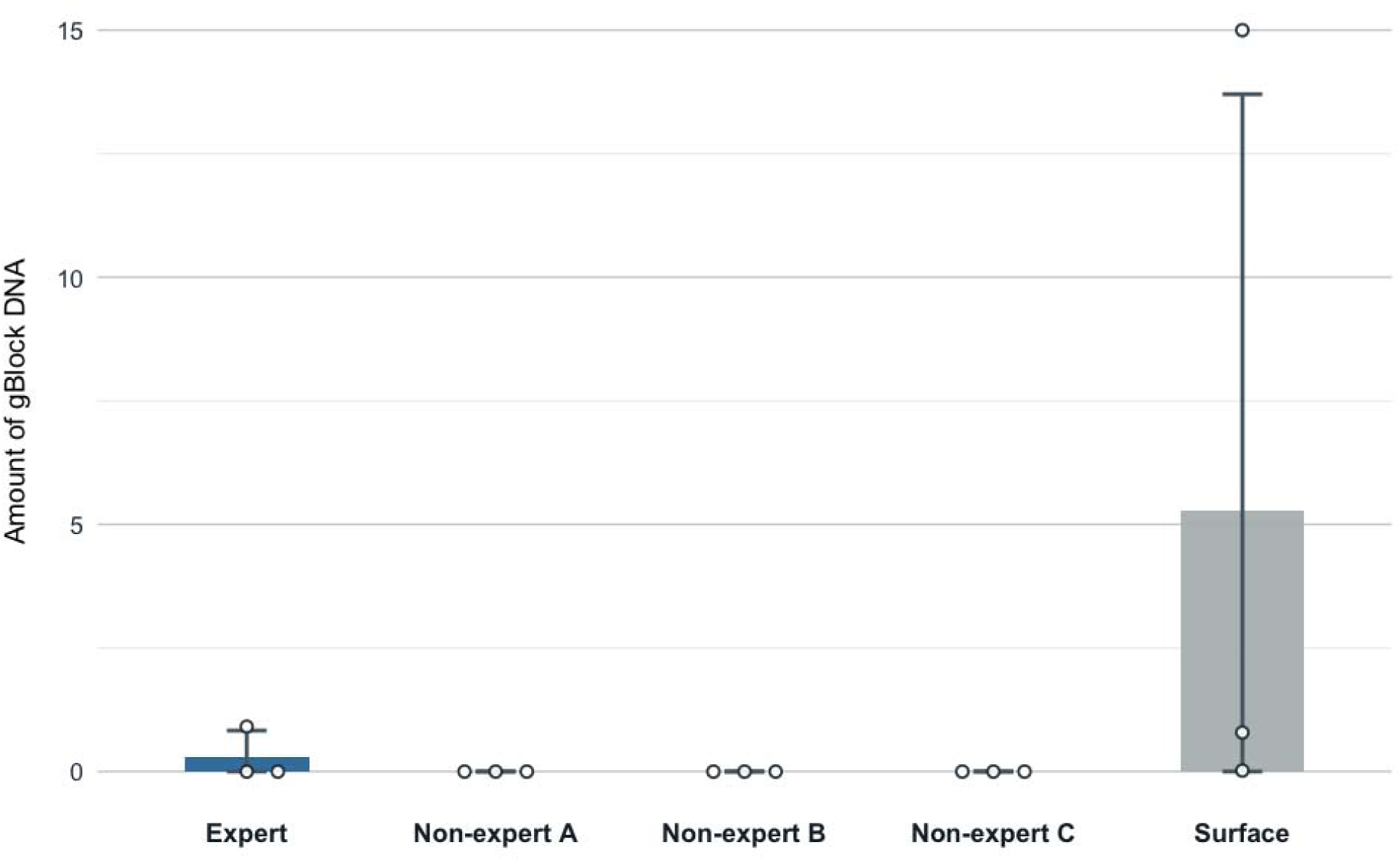
Comparison of the average gBlock DNA molecule per µl between the surface samples and the samples taken by one expert and three non-expert participants.

After the non-experts conducted a clean sediment sampling following the provided protocols, contamination levels in their samples were significantly reduced (Two-tailed Student T-test *p-value* = 0.030). In most cases, gBlock DNA levels dropped to near zero, demonstrating the effectiveness of the cleaning procedures in removing artificially introduced contamination. (Fig. 1). The samples collected by the expert also exhibited minimal contamination except for one sample, which still showed a low level of gBlock DNA, possibly due to accidental contamination from spraying applications. (Fig. 3)

#### Sand-rich soil

In total, fifteen samples were collected: (a) three artificially contaminated surface sediment samples and (b) twelve samples collected by four non-expert participants (three samples each). The surface sediment samples from the sand-rich soil site showed high levels of artificial gBlock DNA contamination, establishing a consistent baseline for evaluating the effectiveness of the cleaning protocols. The cleaning procedures performed by the non-experts significantly reduced contamination levels (Two-tailed Student T-test *p-value* = 0.000) (Fig. 4). However, unlike the clay-rich soil samples, the samples from sand-rich soil exhibited relatively higher residual gBlock DNA levels after a thorough clean sampling.

**Figure 4.**
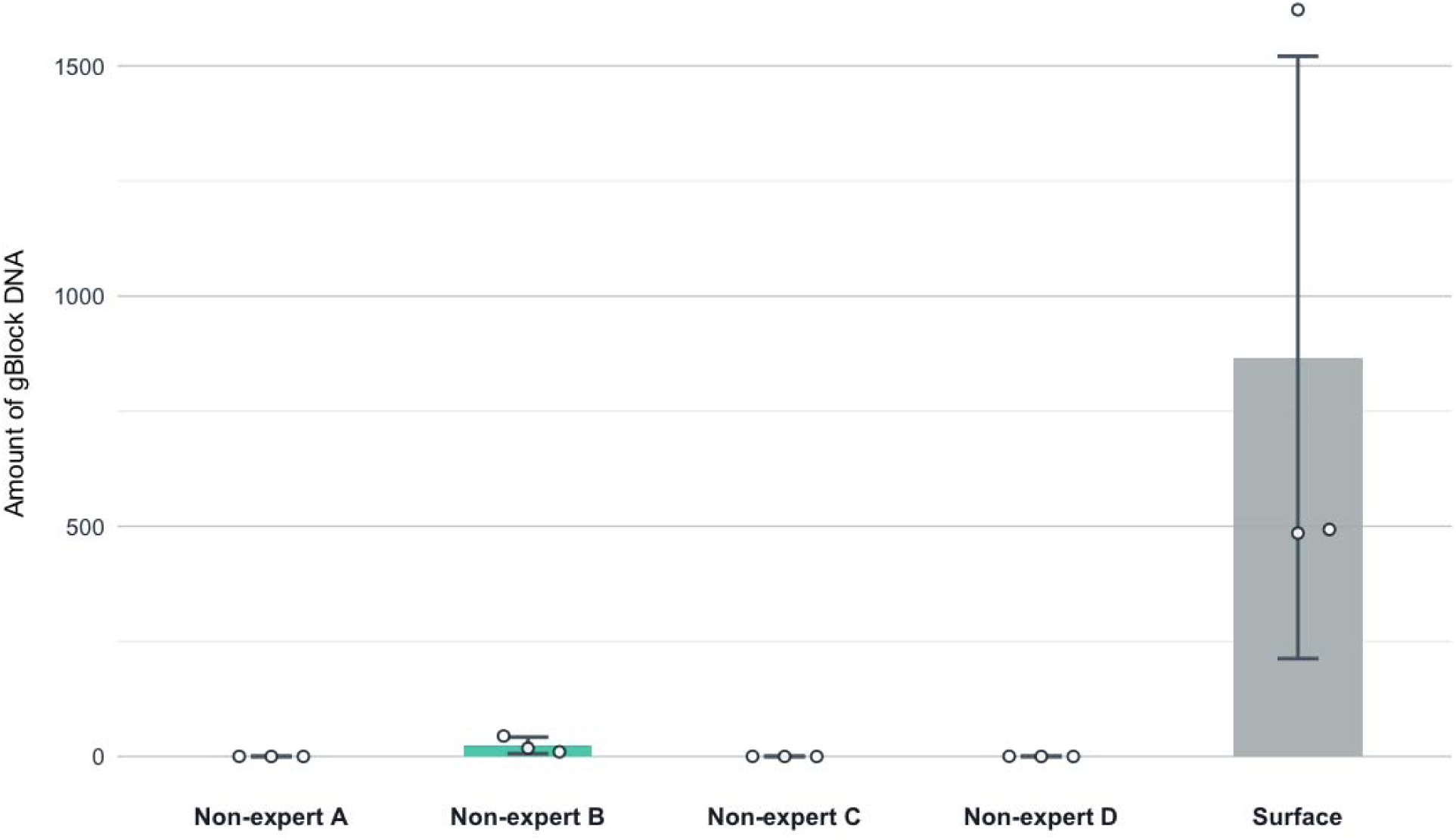
Comparison of the average gBlock DNA molecule per µl between the surface samples and the samples taken by four non-expert participants.

### Sampling from a pollen core

After applying the artificial surface contamination, six subsamples were collected from the pollen core to evaluate the distribution of gBlock DNA contamination across different depths. Two subsamples were taken from the contaminated surface, two from the midpoint using the clean subsampling protocol (Epp et al., 2019), and two from the bottom of the pollen core, which had been in contact with its surface.

*Surface subsamples*: The surface subsamples exhibited significantly high levels of artificial DNA contamination, comparing the samples from the mid-point and the bottom (Two-tailed Student T-test *p-values* = 0.024 and 0.024, respectively). This result aligns with expectations, as these samples directly interacted with the sprayed gBlock DNA during the contamination simulation (Fig. 5).

**Figure 5.**
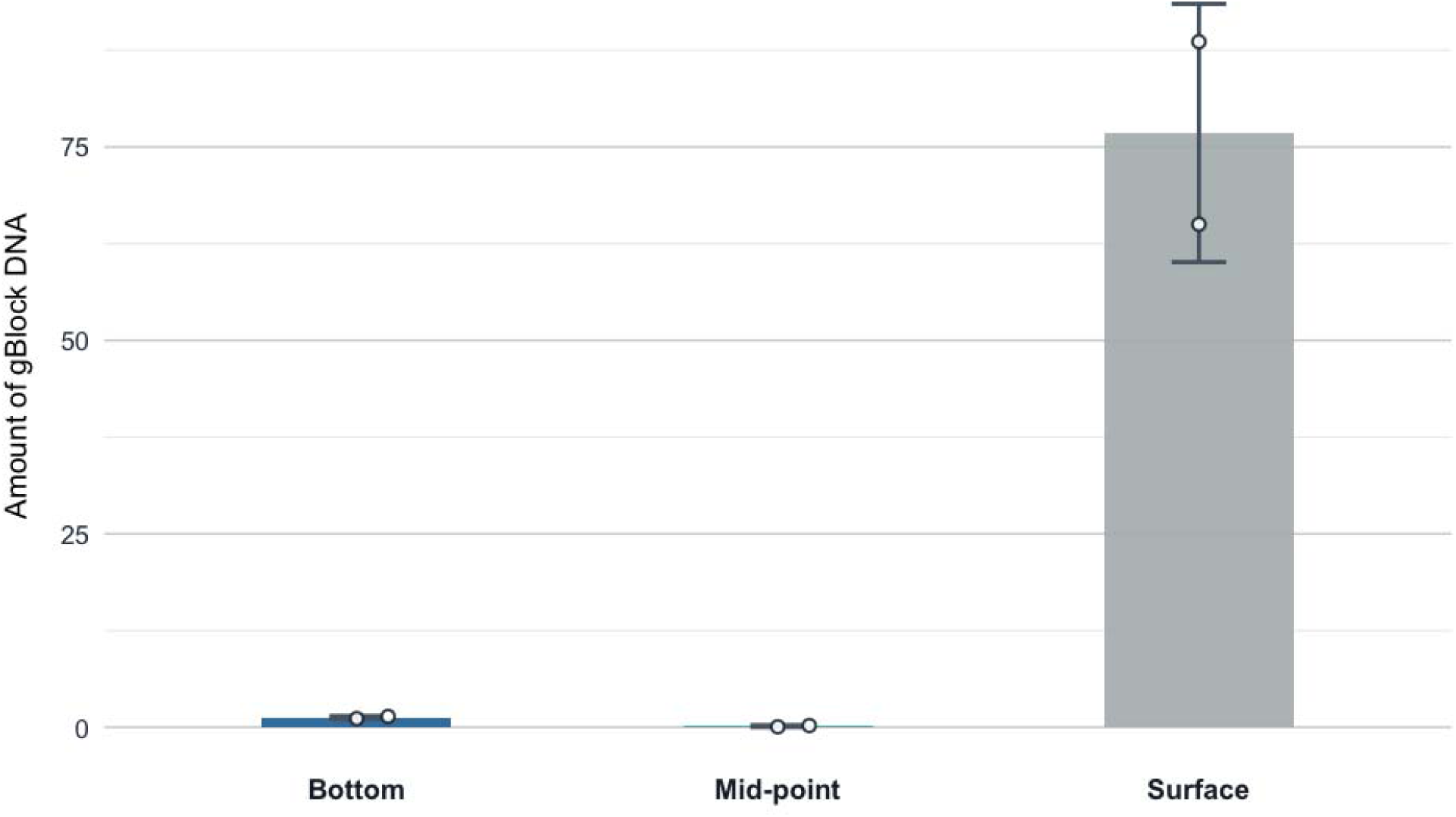
Comparison of the average gBlock DNA molecule per µl between the samples taken from the surface, the mid-point, and the bottom of the pollen core.

*Midpoint subsamples*: The midpoint subsamples showed significantly minimal contamination compared to samples from the bottom of the core (Two-tailed Student T-test *p-value* = 0.021), with only trace amounts of artificial DNA detected. The low contamination levels suggest that the clean sampling protocol effectively minimized contamination from the surface layers, indicating that the mid-point is ideal for collecting sediment samples for sedaDNA studies. However, trace gBlock DNA may indicate minor contamination introduced via airborne particles, as the spraying and subsampling were performed in a closed room (Fig. 5).

*Bottom-point subsamples*: The bottom-point subsamples exhibited slightly higher gBlock DNA amounts than the mid-point subsamples. This increase is likely due to the bottom point’s proximity to the core’s inner surfaces, which had been sprayed with artificial DNA before core sampling. The closeness to these sprayed surfaces may have contributed to a greater likelihood of contamination during sampling (Fig. 5).

## Discussion

This study underscores the significance of careful sediment sampling to minimize surface contamination, a crucial factor that can obscure authentic sedaDNA signals. Regardless of being modern or ancient, surface contamination can skew data interpretation and undermine the reliability of sedaDNA analyses. While some samples exhibited low surface contamination, clean sampling protocols effectively allow bioinformatic tools to filter out residual contamination, thus providing a solid starting point for further analysis (Michelsen et al., 2022; Pochon et al., 2023).

Overall, the results from both soil types demonstrate the effectiveness of clean sampling protocols in significantly reducing artificially introduced gBlock DNA contamination. However, sand-rich soil samples showed higher residual contamination than clay-rich soil samples. This variation highlights the influence of soil composition on contamination dynamics (Freeman et al., 2023). Additionally, the pollen core sampling experiment demonstrated the significance of depth in reducing surface contamination, with the midpoint recognized as the optimal point for sedaDNA subsampling. These findings collectively emphasize the essential role of clean sampling protocols in minimizing the risk of surface contamination.

Our findings also show no statistically significant differences in contamination levels between experts and non-experts. This indicates that when clean sampling protocols are complemented with thorough visual (e.g., video or in-person) and written instructional materials, archaeologists are as well-equipped as specialists to collect sedaDNA samples. At the same time, both groups significantly reduced surface contamination compared to surface samples. Allowing non-experts to conduct clean sampling with minimal supervision has significant implications for archaeological fieldwork. By decreasing dependence on expert oversight, the spatial and temporal resolution of sedaDNA studies can be significantly enhanced, making these analyses more accessible and scalable.

Furthermore, our findings indicate that subsampling sediments with minimal surface contamination from previously collected materials, such as pollen cores, is a viable approach. This highlights the potential of archival sediment collections for sedaDNA analysis, expanding the opportunities for reconstructing past environments and biodiversity (Özdoğan et al., 2024).

However, contamination during sampling remains a significant risk. To reduce the number of steps where contamination could be introduced, we recommend using sterile tools and sampling sediments into sterile tubes at excavation sites, especially for sites with high importance. However, in cases where sterile tools are unavailable, archaeologists can safely use standard pollen cores for sampling. Pollen cores are a safe option for sedaDNA sampling and bring logistic advantages, such as the flexibility for unplanned sampling when needed or the potential of transporting a whole stratigraphic context, which allows more comprehensive sampling. However, we strongly advise conducting core subsampling in an ancient DNA laboratory or a clean room while exercising extreme caution to minimize surface and airborne contamination, as described in appendix A.1.

Based on these findings, our study emphasizes that sedaDNA sampling can be easily integrated into archaeological workflows. Incorporating sedaDNA data has the potential to complement and cross-validate archaeobotanical and archaeozoological findings, offering a more holistic perspective for addressing archaeological questions related to past environments and biodiversity.

## Conclusion

The analyses provide valuable insights into the possible sources of contamination during sediment sampling and handling, highlighting the feasibility of safely using previously collected pollen cores when clean sampling practices are applied. This study also underscores the critical value of clean sampling protocols in making sedaDNA research more accessible. By enabling non-experts to collect and subsample materials, including archival sediments, these methods enhance the scalability and reliability of sedaDNA studies and strengthen their potential impact on archaeological research by integrating aDNA analyses into various workflows.

Our results across different soil types confirm the effectiveness of clean sampling protocols in significantly reducing surface contamination and ensuring the integrity of sedaDNA data. Furthermore, the pollen core sampling experiment highlights the importance of clean sampling and subsampling depth in minimizing contamination, with the midpoint of the core identified as the optimal zone for sedaDNA subsampling.

Future research should prioritize investigating other sources of contamination for both modern and ancient environmental contamination. Addressing these contamination risks will further refine sedaDNA methodologies and expand their applications for reconstructing past environments and biodiversity with even greater precision.

## Acknowledgments

We want to extend our appreciation to Irene Mena’s illustrations of sampling processes (Fig. 1 and 2). We also thank Wouter Vos and Folkert Westra for arranging excavation sites for sampling, as well as Lara Boon, Arjan Ruiter, Ruun Nillesen, Timothy van der Knaap, Maurya Hoppenbrouwers, and Jan Hofsteenge for participating in the sampling.

This paper derives from the project ‘Constructing the Limes: Employing citizen science to understand borders and border systems from the Roman period until today’ (C-Limes), funded by the Dutch Research Council (NWO) as part of the Dutch Research Agenda. (2021–2026, Project number: NWA.1292.19.364)

## Author Contributions

K.T. Özdoğan, G.A. de Groot, and G. Plets designed the study. K.T. Özdoğan conducted the field experiments, wrote the manuscript, and prepared Figures 3-5. A. de Groot and G. Plets made additions and suggestions to the whole manuscript, and both are considered senior authors. F. Furni performed the statistical analysis and made suggestions for the manuscript. I. Laros conducted the lab work and made additions and suggestions on the methods. All authors reviewed and approved the final version of the manuscript.

## Declaration of generative AI and AI-assisted technologies in the writing process

During the preparation of this work, the author used Grammarly and ChatGPT in order to improve language and readability with caution. After using these tools, the authors reviewed and edited the content as needed and take full responsibility for the content of the publication.

## Appendices

## Appendix 1

### Protocols for sedaDNA Sampling

#### Protocol for Tube Sampling from the Archaeological Section

Materials:

- Water-Bleach (5%) solution
- Water
- Bleach
- Plastic bottle
- Disposable gloves
- Paper towels
- Face masks
- Disposable coveralls
- Disposable spatulas (or metal spatulas)
- Sterile spatula
- 50 mL sterile tubes
- Small zipped plastic bags

Equipment Setup & Procedure:

1) Dress up:
  a) Put on the gloves first, and clean the hands with bleach.
  b) Put the mask on.
  c) Put on the hair net and the coverall (optional).
  d) Put on another pair of gloves and clean your hands with bleach.
2) Take a spatula and clean it with bleach.
3) Use it to dig into the section about 3 cm.
4) Take another spatula and clean it with bleach. (it can be used again for other samples after cleaning with bleach.)
5) Take a 5/15 mL tube and position it under the dug section. (The needed amount of soil sample is less than 100 mg per extraction (according to Rohland et al., 2018; Zavala et al.,2021))
6) Fill the tube partly with the soil from the section using a sterile spatula.
7) Close the lid and label the tube. (Proper labeling is crucial. A label should include all the information about the sample’s location, sampling date, and (if known) archeological period. The location information should be based on the excavation’s own system.)
8) Put the tube in a labeled zipped bag.
9) Dispose of the spatula and the pair of gloves. (In case of a limited number of materials, clean them intensively with bleach to reuse them.)
10) For the following sample, return to 1d and repeat the remaining steps.

### Protocol for Core Sampling from the Archaeological Section

#### Step-1: Field Sampling

Materials

- Water-Bleach (5%) solution
- Water
- Bleach
- Plastic bottle
- Disposable gloves
- Paper towels
- Face masks
- Disposable coveralls
- Labelling stickers
- Plastic wraps
- Metal or plastic pollen cores
- Hammer
- A trowel

Equipment Setup & Procedure

1) Dress up:
  a) Put on the gloves first, and clean the hands with bleach.
  b) Put the mask on.
  c) Put on the hair net and the coverall (optional).
  d) Put on another pair of gloves and clean your hands with bleach.
2) Take a spatula and clean it with bleach.
3) Clean the surface with the spatula (or trowel) (It can be used again for other samples after cleaning with bleach.)
4) Take the pollen core and place it in the section with the help of a hammer.
5) Take the core out with the help of a travel.
6) Wrap the core with the plastic wrap.
7) Put the sticker on the core and label the core. (Proper labeling is crucial. A label should include all the information about the sample’s location, sampling date, and (if known) archeological period. The location information should be based on the excavation’s own system.)
8) Dispose of the spatula and the pair of gloves. (In case of a limited number of materials, clean them intensively with bleach to reuse them.)
9) For the following sample, return to 1d and repeat the remaining steps.

#### Step-2: Subsampling

Materials:

- Water-Bleach (5%) solution
- Water
- Bleach
- Plastic bottle
- Disposable gloves
- Paper towels
- Face masks
- Disposable coveralls
- Disposable spatulas (or metal spatulas)
- Sterile spatula
- 50 mL sterile tubes
- Small zipped plastic bags

Equipment Setup & Procedure:

1) Dress up:
  a) Put on the gloves first, and clean the hands with bleach.
  b) Put the mask on.
  c) Put on the hair net and the coverall (optional).
  d) Put on another pair of gloves and clean your hands with bleach.
2) Take a spatula and clean it with bleach.
3) Clean the surface of the pollen core with the spatula (It can be used again for other samples after cleaning with bleach.)
4) Dig 2-5cm (close to the midpoint) through the cleaned area with a sterile spatula.
5) Fill up the sterile tube with soil from the midpoint of the core with a sterile spatula.
6) Close the lid and label the tube. (Proper labeling is crucial. A label should include all the information about the sample’s location, sampling date, and (if known) archeological period. The location information should be based on the excavation’s own system.)
7) Put the tube in a labeled zipped bag.
8) Dispose of the spatula and the pair of gloves. (In case of a limited number of materials, clean them intensively with bleach to reuse them.)
9) For the following sample, return to 1d and repeat the remaining steps.

Protocols and an instruction video is available on GitHub: https://github.com/ktozdogan/sedaDNAsampling

## Appendix 2

Based on the comparisons of different concentrations (1e+4, 1e+6, 1e+8), it was determined that 1e+7 c/µl is the optimal detectable concentration and was used in the field experiment.

1ml solution in a 2ml sprayer:

- 90µl Buffer AE,
- 10µl 1e+9 gBlock DNA
- 900µl technical ethanol.

The 2ml spraying bottles have a spray volume of ∼50µl. All the sampling locations were sprayed twice in order to reach a detectable level of gBlock concentration.

The ddPCR was executed on a BioRad QX200 Droplet Digital PCR System, with the emulsion created on a BioRad QX100 droplet generator. The fluorescent signal of the tracer DNA was measured in the VIC channel, and the FAM channel was used for an internal positive control (IPC) TCCB.

The ddPCR reaction consisted of 1x ddPCR Supermix for Probes (BioRad), 900nM Tracer-F, 900nM Tracer-R, 250nM Tracer-probe, 450nM TCCBL, 450nM TCCBR and 250nM TCCB-probe, approximately 50 copies of TCCB positive control DNA and 4µl of sedaDNA extract.

The PCR’s temperature program consisted of a ten-minute hold at 95°C, followed by 40 cycles at 94°C for 30 seconds and 60°C for 1 minute, an enzyme deactivation at 98°C for 10 minutes, and finally, an infinite hold at 4°C.

Complete Tracer GBlock DNA sequence:

TATTTTAGTGGTCATGGGTTTTGGTCCGCCCGAGCGGTGCAACCGATTAGGACCAT GTAAAACATTTGTTACAAGTCTTCTTTTAAACACAATCTTCCTGCTCAGTGGCGCAT GATTATCGTTGTTGCTAGCCAGCGTGGTAAGTAACAGCACCACTGCGAGCCTAATG TGCCCTTTCCACGAACACAGGGCTGTCCGATCCTATATTAGGACTCCGCAATGGG GTTAGCAAGTCGCACCCTAAACGATGTTGAAGACTCGCGATGTACATGCTCTGGTA CAATACATACGTGTTCCGGCAAGCGTGATATTGCTCTTTCGTATAGTTACCATGGC AATGCTTAGAACAATACTAATGTTGTAATCTGTCGCTATGTTAAGAACCGCACGAAC CACAGAGCATAAAGAGAACCTCTAGCTCCTTTACAAAGTACTGGTTCCCTTTCCAG CGGGATGCCTTATCTAAACGCAATGACAGACGTATTCCTCAGGCCACATCGCTTCC TACTTCCGCTGGGATCCATCATTGGCGGCCGAAGCCGCCATTCCATAGTGAGTCC TTCGTCTGTGTCTTTCTGTGCCAGATCGTCTAGCAAATTGCCGATCCAGTTTATCTC ACGAAACTATAGTCGTACAGACCGAAATCTTAAGTCAAATCACGCGACTAGGCTCA GCTC

The underlined part is the sequence amplified by the used primers. In bold are the primer sequences.

The TCCB positive DNA control is a quantified PCR product of *Triturus cristatus* tissue amplified by the TCCBL and TCCBR primers (Thomsen et al., 2012).

Primers used

Target

Tracer-F: AAG CGT GAT ATT GCT CTT TCG TAT AG

Tracer-R: ACA TAG CGA CAG ATT ACA ACA TTA GTA TTG

Tracer-probe: VIC-TAC CAT GGC AAT GCT-MGB-NFQ

IPC

TCCBL: CGTAAACTACGGCTGACTAGTACGAA

TCCBR: CCGATGTGTATGTAGATGCAAACA

TCCB-probe: 6FAM-CATCCACGCTAACGGAGCCTCGC-MGB-NFQ

## References

1. Aldeias, V., & Stahlschmidt, M. C. (2024). Sediment DNA can revolutionize archaeology—If it is used the right way. Proceedings of the National Academy of Sciences, 121(26), e2317042121. 10.1073/pnas.2317042121

2. Alsos, I. G., Boussange, V., Rijal, D. P., Beaulieu, M., Brown, A. G., Herzschuh, U., Svenning, J.-C., & Pellissier, L. (2024). Using ancient sedimentary DNA to forecast ecosystem trajectories under climate change. Philosophical Transactions of the Royal Society B: Biological Sciences, 379(1902), 20230017. 10.1098/rstb.2023.0017

3. Andersen, K., Bird, K. L., Rasmussen, M., Haile, J., Breuning-Madsen, H., Kjær, K. H., Orlando, L., Gilbert, M. T. P., & Willerslev, E. (2012a). Meta-barcoding of ‘dirt’ DNA from soil reflects vertebrate biodiversity. Molecular Ecology, 21(8), 1966–1979. 10.1111/j.1365-294X.2011.05261.x

4. Andersen, K., Bird, K. L., Rasmussen, M., Haile, J., Breuning-Madsen, H., Kjær, K. H., Orlando, L., Gilbert, M. T. P., & Willerslev, E. (2012b). Meta-barcoding of ‘dirt’ DNA from soil reflects vertebrate biodiversity. Molecular Ecology, 21(8), 1966–1979. 10.1111/j.1365-294X.2011.05261.x

5. Ariza, M., Fouks, B., Mauvisseau, Q., Halvorsen, R., Alsos, I. G., & de Boer, H. J. (2023). Plant biodiversity assessment through soil eDNA reflects temporal and local diversity. Methods in Ecology and Evolution, 14(2), 415–430. 10.1111/2041-210X.13865

6. Armbrecht, L. H., Coolen, M. J. L., Lejzerowicz, F., George, S. C., Negandhi, K., Suzuki, Y., Young, J., Foster, N. R., Armand, L. K., Cooper, A., Ostrowski, M., Focardi, A., Stat, M., Moreau, J. W., & Weyrich, L. S. (2019). Ancient DNA from marine sediments: Precautions and considerations for seafloor coring, sample handling and data generation. Earth-Science Reviews, 196, 102887. 10.1016/j.earscirev.2019.102887

7. Armbrecht, L., Weber, M. E., Raymo, M. E., Peck, V. L., Williams, T., Warnock, J., Kato, Y., Hernández-Almeida, I., Hoem, F., Reilly, B., Hemming, S., Bailey, I., Martos, Y. M., Gutjahr, M., Percuoco, V., Allen, C., Brachfeld, S., Cardillo, F. G., Du, Z., … Zheng, X. (2022). Ancient marine sediment DNA reveals diatom transition in Antarctica. Nature Communications, 13(1), 5787. 10.1038/s41467-022-33494-4

8. Boivin, S., Bourceret, A., Maurice, K., Laurent-Webb, L., Figura, T., Bourillon, J., Nespoulous, J., Domergue, O., Chaintreuil, C., Boukcim, H., Selosse, M.-A., Fiema, Z., Botte, E., Nehme, L., & Ducousso, M. (2024). Revealing human impact on natural ecosystems through soil bacterial DNA sampled from an archaeological site. Environmental Microbiology, 26(1), e16546. 10.1111/1462-2920.16546

9. Borry, M., Cordova, B., Perri, A., Wibowo, M., Honap, T. P., Ko, J., Yu, J., Britton, K., Girdland-Flink, L., Power, R. C., Stuijts, I., Salazar-García, D. C., Hofman, C., Hagan, R., Kagoné, T. S., Meda, N., Carabin, H., Jacobson, D., Reinhard, K., … Warinner, C. (2020). CoproID predicts the source of coprolites and paleofeces using microbiome composition and host DNA content. PeerJ, 8, e9001. 10.7717/peerj.9001

10. Braadbaart, F., Reidsma, F. H., Roebroeks, W., Chiotti, L., Slon, V., Meyer, M., Théry-Parisot, I., van Hoesel, A., Nierop, K. G. J., Kaal, J., van Os, B., & Marquer, L. (2020). Heating histories and taphonomy of ancient fireplaces: A multi-proxy case study from the Upper Palaeolithic sequence of Abri Pataud (Les Eyzies-de-Tayac, France). Journal of Archaeological Science: Reports, 33, 102468. 10.1016/j.jasrep.2020.102468

11. Conte, J., Potoczniak, M. J., & Tobe, S. S. (2018). Using synthetic oligonucleotides as standards in probe-based qPCR. BioTechniques, 64(4), 177–179. 10.2144/btn-2018-2000

12. Crump, S. E., Fréchette, B., Power, M., Cutler, S., de Wet, G., Raynolds, M. K., Raberg, J. H., Briner, J. P., Thomas, E. K., Sepúlveda, J., Shapiro, B., Bunce, M., & Miller, G. H. (2021). Ancient plant DNA reveals High Arctic greening during the Last Interglacial. Proceedings of the National Academy of Sciences, 118(13), e2019069118. 10.1073/pnas.2019069118

13. Dabney, J., Meyer, M., & Pääbo, S. (2013). Ancient DNA Damage. Cold Spring Harbor Perspectives in Biology, 5(7), a012567. 10.1101/cshperspect.a012567

14. DNeasy PowerSoil Pro Kit Handbook—QIAGEN. (n.d.). Retrieved November 24, 2024, from https://www.qiagen.com/us/resources/resourcedetail?id=9bb59b74-e493-4aeb-b6c1-f660852e8d97&lang=en

15. Dommain, R., Andama, M., McDonough, M. M., Prado, N. A., Goldhammer, T., Potts, R., Maldonado, J. E., Nkurunungi, J. B., & Campana, M. G. (2020). The Challenges of Reconstructing Tropical Biodiversity With Sedimentary Ancient DNA: A 2200-Year-Long Metagenomic Record From Bwindi Impenetrable Forest, Uganda. Frontiers in Ecology and Evolution, 8. 10.3389/fevo.2020.00218

16. Edwards, M. E., Alsos, I. G., Yoccoz, N., Coissac, E., Goslar, T., Gielly, L., Haile, J., Langdon, C. T., Tribsch, A., Binney, H. A., Stedingk, H. von, & Taberlet, P. (2018). Metabarcoding of modern soil DNA gives a highly local vegetation signal in Svalbard tundra. The Holocene. 10.1177/0959683618798095

17. Epp, L. S., Zimmermann, H. H., & Stoof-Leichsenring, K. R. (2019). Sampling and Extraction of Ancient DNA from Sediments. In B. Shapiro, A. Barlow, P. D. Heintzman, M. Hofreiter, J. L. A. Paijmans, & A. E. R. Soares (Eds.), Ancient DNA: Methods and Protocols (pp. 31–44). Springer. 10.1007/978-1-4939-9176-1_5

18. Essel, E., Zavala, E. I., Schulz-Kornas, E., Kozlikin, M. B., Fewlass, H., Vernot, B., Shunkov, M. V., Derevianko, A. P., Douka, K., Barnes, I., Soulier, M.-C., Schmidt, A., Szymanski, M., Tsanova, T., Sirakov, N., Endarova, E., McPherron, S. P., Hublin, J.-J., Kelso, J., … Meyer, M. (2023). Ancient human DNA recovered from a Palaeolithic pendant. Nature, 618(7964), 328–332. 10.1038/s41586-023-06035-2

19. Foley, B. P., Hansson, M. C., Kourkoumelis, D. P., & Theodoulou, T. A. (2012). Aspects of ancient Greek trade re-evaluated with amphora DNA evidence. Journal of Archaeological Science, 39(2), 389–398. 10.1016/j.jas.2011.09.025

20. Freeman, C. L., Dieudonné, L., Agbaje, O. B. A., Žure, M., Sanz, J. Q., Collins, M., & Sand, K. K. (2023). Survival of environmental DNA in sediments: Mineralogic control on DNA taphonomy. Environmental DNA, 5(6), 1691–1705. 10.1002/edn3.482

21. Fulton, T. L., & Shapiro, B. (2019). Setting Up an Ancient DNA Laboratory. In B. Shapiro, A. Barlow, P. D. Heintzman, M. Hofreiter, J. L. A. Paijmans, & A. E. R. Soares (Eds.), Ancient DNA: Methods and Protocols (pp. 1–13). Springer. 10.1007/978-1-4939-9176-1_1

22. Furtwängler, A., Reiter, E., Neumann, G. U., Siebke, I., Steuri, N., Hafner, A., Lösch, S., Anthes, N., Schuenemann, V. J., & Krause, J. (2018). Ratio of mitochondrial to nuclear DNA affects contamination estimates in ancient DNA analysis. Scientific Reports, 8(1), 14075. 10.1038/s41598-018-32083-0

23. Gelabert, P., Sawyer, S., Bergström, A., Margaryan, A., Collin, T. C., Meshveliani, T., Belfer-Cohen, A., Lordkipanidze, D., Jakeli, N., Matskevich, Z., Bar-Oz, G., Fernandes, D. M., Cheronet, O., Özdoğan, K. T., Oberreiter, V., Feeney, R. N. M., Stahlschmidt, M. C., Skoglund, P., & Pinhasi, R. (2021). Genome-scale sequencing and analysis of human, wolf, and bison DNA from 25,000-year-old sediment. Current Biology, 31(16), 3564–3574.e9. 10.1016/j.cub.2021.06.023

24. Giguet-Covex, C., Jelavić, S., Foucher, A., Morlock, M. A., Wood, S. A., Augustijns, F., Domaizon, I., Gielly, L., & Capo, E. (2023). The Sources and Fates of Lake Sedimentary DNA. In E. Capo, C. Barouillet, & J. P. Smol (Eds.), Tracking Environmental Change Using Lake Sediments: Volume 6: Sedimentary DNA (pp. 9–52). Springer International Publishing. 10.1007/978-3-031-43799-1_2

25. Green, E. J., & Speller, C. F. (2017). Novel Substrates as Sources of Ancient DNA: Prospects and Hurdles. Genes, 8(7), Article 7. 10.3390/genes8070180

26. Hebda, C. F. G., McLaren, D., Mackie, Q., Fedje, D., Pedersen, M. W., Willerslev, E., Brown, K. J., & Hebda, R. J. (2022). Late Pleistocene palaeoenvironments and a possible glacial refugium on northern Vancouver Island, Canada: Evidence for the viability of early human settlement on the northwest coast of North America. Quaternary Science Reviews, 279, 107388. 10.1016/j.quascirev.2022.107388

27. Hunter, M. E., Dorazio, R. M., Butterfield, J. S. S., Meigs-Friend, G., Nico, L. G., & Ferrante, J. A. (2017). Detection limits of quantitative and digital PCR assays and their influence in presence–absence surveys of environmental DNA. Molecular Ecology Resources, 17(2), 221–229. 10.1111/1755-0998.12619

28. Kavlick, M. F. (2018). Development of a Universal Internal Positive Control. BioTechniques, 65(5), 275–280. 10.2144/btn-2018-0034

29. Kjær, K. H., Winther Pedersen, M., De Sanctis, B., De Cahsan, B., Korneliussen, T. S., Michelsen, C. S., Sand, K. K., Jelavić, S., Ruter, A. H., Schmidt, A. M. A., Kjeldsen, K. K., Tesakov, A. S., Snowball, I., Gosse, J. C., Alsos, I. G., Wang, Y., Dockter, C., Rasmussen, M., Jørgensen, M. E., … Willerslev, E. (2022). A 2-million-year-old ecosystem in Greenland uncovered by environmental DNA. Nature, 612(7939), 283–291. 10.1038/s41586-022-05453-y

30. Knapp, M., Clarke, A. C., Horsburgh, K. A., & Matisoo-Smith, E. A. (2012). Setting the stage – Building and working in an ancient DNA laboratory. Annals of Anatomy – Anatomischer Anzeiger, 194(1), 3–6. 10.1016/j.aanat.2011.03.008

31. Llamas, B., Valverde, G., Fehren-Schmitz, L., Weyrich, L. S., Cooper, A., & Haak, W. (2017). From the field to the laboratory: Controlling DNA contamination in human ancient DNA research in the high-throughput sequencing era. STAR: Science & Technology of Archaeological Research, 3(1), 1–14. 10.1080/20548923.2016.1258824

32. Michelsen, C., Pedersen, M. W., Fernandez-Guerra, A., Zhao, L., Petersen, T. C., & Korneliussen, T. S. (2022). metaDMG – A Fast and Accurate Ancient DNA Damage Toolkit for Metagenomic Data (p. 2022.12.06.519264). bioRxiv. 10.1101/2022.12.06.519264

33. Murchie, T. J., Karpinski, E., Eaton, K., Duggan, A. T., Baleka, S., Zazula, G., MacPhee, R. D. E., Froese, D., & Poinar, H. N. (2022). Pleistocene mitogenomes reconstructed from the environmental DNA of permafrost sediments. Current Biology, 32(4), 851–860.e7. 10.1016/j.cub.2021.12.023

34. Murchie, T. J., Monteath, A. J., Mahony, M. E., Long, G. S., Cocker, S., Sadoway, T., Karpinski, E., Zazula, G., MacPhee, R. D. E., Froese, D., & Poinar, H. N. (2021). Collapse of the mammoth-steppe in central Yukon as revealed by ancient environmental DNA. Nature Communications, 12(1), 7120. 10.1038/s41467-021-27439-6

35. Nguyen, N.-L., Devendra, D., Szymańska, N., Greco, M., Angeles, I. B., Weiner, A. K. M., Ray, J. L., Cordier, T., De Schepper, S., Pawłowski, J., & Pawłowska, J. (2023). Sedimentary ancient DNA: A new paleogenomic tool for reconstructing the history of marine ecosystems. Frontiers in Marine Science, 10. 10.3389/fmars.2023.1185435

36. Orlando, L., Allaby, R., Skoglund, P., Der Sarkissian, C., Stockhammer, P. W., Ávila-Arcos, M. C., Fu, Q., Krause, J., Willerslev, E., Stone, A. C., & Warinner, C. (2021). Ancient DNA analysis. Nature Reviews Methods Primers, 1(1), Article 1. 10.1038/s43586-020-00011-0

37. Özdoğan, K. T., Gelabert, P., Hammers, N., Altınışık, N. E., de Groot, A., & Plets, G. (2024). Archaeology meets environmental genomics: Implementing sedaDNA in the study of the human past. Archaeological and Anthropological Sciences, 16(7), 108. 10.1007/s12520-024-01999-2

38. Pedersen, M. W., Sanctis, B. D., Saremi, N. F., Sikora, M., Puckett, E. E., Gu, Z., Moon, K. L., Kapp, J. D., Vinner, L., Vardanyan, Z., Ardelean, C. F., Arroyo-Cabrales, J., Cahill, J. A., Heintzman, P. D., Zazula, G., MacPhee, R. D. E., Shapiro, B., Durbin, R., & Willerslev, E. (2021). Environmental genomics of Late Pleistocene black bears and giant short-faced bears. Current Biology, 31(12), 2728–2736.e8. 10.1016/j.cub.2021.04.027

39. Peyrégne, S., & Peter, B. M. (2020). AuthentiCT: A model of ancient DNA damage to estimate the proportion of present-day DNA contamination. Genome Biology, 21(1), 246. 10.1186/s13059-020-02123-y

40. Peyrégne, S., & Prüfer, K. (2020). Present-Day DNA Contamination in Ancient DNA Datasets. BioEssays, 42(9), 2000081. 10.1002/bies.202000081

41. Pochon, Z., Bergfeldt, N., Kırdök, E., Vicente, M., Naidoo, T., van der Valk, T., Altınışık, N. E., Krzewińska, M., Dalén, L., Götherström, A., Mirabello, C., Unneberg, P., & Oskolkov, N. (2023). aMeta: An accurate and memory-efficient ancient metagenomic profiling workflow. Genome Biology, 24(1), 242. 10.1186/s13059-023-03083-9

42. Renaud, G., Slon, V., Duggan, A. T., & Kelso, J. (2015). Schmutzi: Estimation of contamination and endogenous mitochondrial consensus calling for ancient DNA. Genome Biology, 16(1), 224. 10.1186/s13059-015-0776-0

43. Sabin, S., Yeh, H.-Y., Pluskowski, A., Clamer, C., Mitchell, P. D., & Bos, K. I. (2020). Estimating molecular preservation of the intestinal microbiome via metagenomic analyses of latrine sediments from two medieval cities. Philosophical Transactions of the Royal Society B. 10.1098/rstb.2019.0576

44. Seeber, P. A., Batke, L., Dvornikov, Y., Schmidt, A., Wang, Y., Stoof-Leichsenring, K., Moon, K., Vohr, S. H., Shapiro, B., & Epp, L. S. (2024). Mitochondrial genomes of Pleistocene megafauna retrieved from recent sediment layers of two Siberian lakes. eLife, 12, RP89992. 10.7554/eLife.89992

45. Seersholm, F. V., Pedersen, M. W., Søe, M. J., Shokry, H., Mak, S. S. T., Ruter, A., Raghavan, M., Fitzhugh, W., Kjær, K. H., Willerslev, E., Meldgaard, M., Kapel, C. M. O., & Hansen, A. J. (2016). DNA evidence of bowhead whale exploitation by Greenlandic Paleo-Inuit 4,000 years ago. Nature Communications, 7(1), 13389. 10.1038/ncomms13389

46. Selway, C. A., Armbrecht, L., & Thornalley, D. (2022). An Outlook for the Acquisition of Marine Sedimentary Ancient DNA (sedaDNA) From North Atlantic Ocean Archive Material. Paleoceanography and Paleoclimatology, 37(5), e2021PA004372. 10.1029/2021PA004372

47. Skoglund, P., Northoff, B. H., Shunkov, M. V., Derevianko, A. P., Pääbo, S., Krause, J., & Jakobsson, M. (2014). Separating endogenous ancient DNA from modern day contamination in a Siberian Neandertal. Proceedings of the National Academy of Sciences, 111(6), 2229–2234. 10.1073/pnas.1318934111

48. Søe, M. J., Nejsum, P., Seersholm, F. V., Fredensborg, B. L., Habraken, R., Haase, K., Hald, M. M., Simonsen, R., Højlund, F., Blanke, L., Merkyte, I., Willerslev, E., & Kapel, C. M. O. (2018). Ancient DNA from latrines in Northern Europe and the Middle East (500 BC– 1700 AD) reveals past parasites and diet. PLOS ONE, 13(4), e0195481. 10.1371/journal.pone.0195481

49. Swango, K. L., Timken, M. D., Chong, M. D., & Buoncristiani, M. R. (2006). A quantitative PCR assay for the assessment of DNA degradation in forensic samples. Forensic Science International, 158(1), 14–26. 10.1016/j.forsciint.2005.04.034

50. Taylor, S. C., Laperriere, G., & Germain, H. (2017). Droplet Digital PCR versus qPCR for gene expression analysis with low abundant targets: From variable nonsense to publication quality data. Scientific Reports, 7(1), 2409. 10.1038/s41598-017-02217-x

51. Thomsen, P. F., Kielgast, J., Iversen, L. L., Wiuf, C., Rasmussen, M., Gilbert, M. T. P., Orlando, L., & Willerslev, E. (2012). Monitoring endangered freshwater biodiversity using environmental DNA. Molecular Ecology, 21(11), 2565–2573. 10.1111/j.1365-294X.2011.05418.x

52. Vernot, B., Zavala, E. I., Gómez-Olivencia, A., Jacobs, Z., Slon, V., Mafessoni, F., Romagné, F., Pearson, A., Petr, M., Sala, N., Pablos, A., Aranburu, A., de Castro, J. M. B., Carbonell, E., Li, B., Krajcarz, M. T., Krivoshapkin, A. I., Kolobova, K. A., Kozlikin, M. B., … Meyer, M. (2021). Unearthing Neanderthal population history using nuclear and mitochondrial DNA from cave sediments. Science, 372(6542), eabf1667. 10.1126/science.abf1667

53. Wang, Y., Pedersen, M. W., Alsos, I. G., De Sanctis, B., Racimo, F., Prohaska, A., Coissac, E., Owens, H. L., Merkel, M. K. F., Fernandez-Guerra, A., Rouillard, A., Lammers, Y., Alberti, A., Denoeud, F., Money, D., Ruter, A. H., McColl, H., Larsen, N. K., Cherezova, A. A., … Willerslev, E. (2021). Late Quaternary dynamics of Arctic biota from ancient environmental genomics. Nature, 600(7887), Article 7887. 10.1038/s41586-021-04016-x

54. Wood, S. A., Pochon, X., Laroche, O., von Ammon, U., Adamson, J., & Zaiko, A. (2019). A comparison of droplet digital polymerase chain reaction (PCR), quantitative PCR and metabarcoding for species-specific detection in environmental DNA. Molecular Ecology Resources, 19(6), 1407–1419. 10.1111/1755-0998.13055

55. Zavala, E. I., Jacobs, Z., Vernot, B., Shunkov, M. V., Kozlikin, M. B., Derevianko, A. P., Essel, E., de Fillipo, C., Nagel, S., Richter, J., Romagné, F., Schmidt, A., Li, B., O’Gorman, K., Slon, V., Kelso, J., Pääbo, S., Roberts, R. G., & Meyer, M. (2021). Pleistocene sediment DNA reveals hominin and faunal turnovers at Denisova Cave. Nature, 595(7867), 399–403. 10.1038/s41586-021-03675-0

56. Zimmermann, H. H., Stoof-Leichsenring, K. R., Kruse, S., Nürnberg, D., Tiedemann, R., & Herzschuh, U. (2021). Sedimentary Ancient DNA From the Subarctic North Pacific: How Sea Ice, Salinity, and Insolation Dynamics Have Shaped Diatom Composition and Richness Over the Past 20,000 Years. Paleoceanography and Paleoclimatology, 36(4), e2020PA004091. 10.1029/2020PA004091

